# CanIsoNet: A Database to Study the Functional Impact of Isoform Switching Events in Diseases

**DOI:** 10.1101/2021.09.17.460795

**Authors:** Tülay Karakulak, Damian Szklarczyk, Cemil Can Saylan, Holger Moch, Christian von Mering, Abdullah Kahraman

## Abstract

**Motivation:** Alternative splicing, as an essential regulatory mechanism in normal mammalian cells, is frequently disturbed in cancer and other diseases. Switches in the expression of most dominant alternative isoforms can alter protein interaction networks of associated genes giving rise to a disease or disease progression. Here, we present CanIsoNet (disease-specific isoform interaction network), a database to view, browse and search these isoform switching events. CanIsoNet is the first webserver that incorporates isoform expression data with STRING interaction networks and ClinVar annotations to predict the pathogenic impact of isoform switching events in various diseases.

**Results:** Data in CanIsoNet can be browsed by disease or searched by genes or isoforms in annotation-rich data tables. Various annotations for 11,072 isoforms and 13,531 unique isoform switching events across 28 different tissue types are provided. The network density score for each disease-specific isoform, PFAM domain IDs of disrupted interactions, domain structure visualization of transcripts and expression data of switched isoforms for each sample are given. Additionally, the genes annotated in ClinVar are highlighted in the interactive interaction network.

**Availability:** CanIsoNet is freely available at https://caniso.net. The source codes can be found under a Creative Common License at https://github.com/kahramanlab/CanIsoNet_Web.

**Supplementary Information:** Supplementary data are available at Bioinformatics online.

## 1 Introduction

Alternative splicing is an essential mechanism to regulate the generation of various mature mRNA transcripts from a single gene^1^. Dysregulation of this mechanism can lead to the overexpression and downregulation of alternative and canonical isoforms, respectively, causing isoform switching events (Climente-González *et al.*, 2017; Kahraman *et al.*, 2020; Vitting-Seerup and Sandelin, 2017). Depending on the length and composition of alternative isoforms, switching event can lead to the loss or gain of interaction domains in affected gene, with functional consequences (Climente-González *et al.*, 2017; Kahraman *et al.*, 2020; Vitting-Seerup and Sandelin, 2017). Various software tools have been implemented to probe these switching events and their functional impact. For example, the IsoformSwitchAnalyzeR software can probe isoform switching events in RNA-seq data and report the functional loss or gain of protein domains (Vitting-Seerup and Sandelin, 2019). More recently, the DIGGER database was introduced as a tool to study the interaction network of protein isoforms and protein domains at an isoform and exon level (Louadi *et al.*, 2021). However, there is no current database or webserver that provides disease-specific isoform switching data for diverse disease types. Here, we describe *CanIsoNet,* a database that merges disease-specific isoform data with STRING (Szklarczyk *et al.*, 2015) protein interaction networks, and pathogenic information from ClinVar to visualize and identify functional impact of isoform switching events in various diseases.

## 2 Implementation

A human and mouse isoform interaction network was constructed based on STRING v11.0 (Kahraman et al., 2020 and Supplementary File 1). The switches in most dominant isoforms were computed and integrated into the isoform interaction network. CanIsoNet was implemented using Python Flask (Python version 3.7) with JavaScript/AJAX extensions, a Bootstrap version 4.3.1 front-end framework and an underlying MySQL database. The Python Plotly library was used for plotting. All plots are zoomable and downloadable. The STRING network was constructed using the STRING API and graphically adjusted to visualise interaction disruptions. Additionally, we integrated disease related genes from ClinVar (data downloaded 13th of July, 2022) to the network visualization via JavaScript. Domain structures of transcripts were generated by the wiggleplotr R package (Kaur Alasoo, 2020). Anatograms were produced by the gganatogram Rshiny app (Maag, 2018). CanIsoNet can be searched or browsed by disease, gene, or isoform names. Files including isoform interaction networks, STRING interaction density scores, or interaction disruption in dominant transcripts for both human and mouse isoforms are available for download from https://caniso.net.

## 3 Results

CanIsoNet currently stores splicing information for a total of 7,172 genes, 11,072 isoforms, and 13,531 unique isoform switching events across 27 cancer types and one mouse trigeminal ganglia sensory tissue. For each disease type, we list all detected disease specific Most Dominant Transcripts (dMDT) and highlight the top 10 most frequent dMDT. The most frequent dMDT are potential diagnostic biomarker candidates (Fig. 1A). CanIsoNet pages dedicated to dMDT, show an isoform-specific interaction network where disruptions are featured via STRING interaction networks (Fig. 1B). Each specific isoform page provides the percentage of disrupted STRING interactions (Fig 1C, right panel), Pfam domain-domain interactions that are lost due to isoform switching events, and a list of lost interactions to disease related genes. Additionally, the network density score, which indicates the number of interactions within the neighbourhood of a protein, is provided (Fig. 1C, left panel, calculated as described in Kahraman et al., 2020). The integration of health and disease related genes from ClinVar (Landrum *et al.*, 2014) is a unique feature of CanIsoNet, which allows users to assess the pathogenicity of a dMDT (Fig 1B, depicted as blue on STRING network). Furthermore, the relative expression values of each dMDT and the median expression value of MDTs in matched normal tissue samples are shown on a sample-specific page for each disease sample (Fig. 1D). In addition, the exon-intron structures of MDTs and dMDTs for each sample are provided (Fig. 1E).

**Figure 1:**
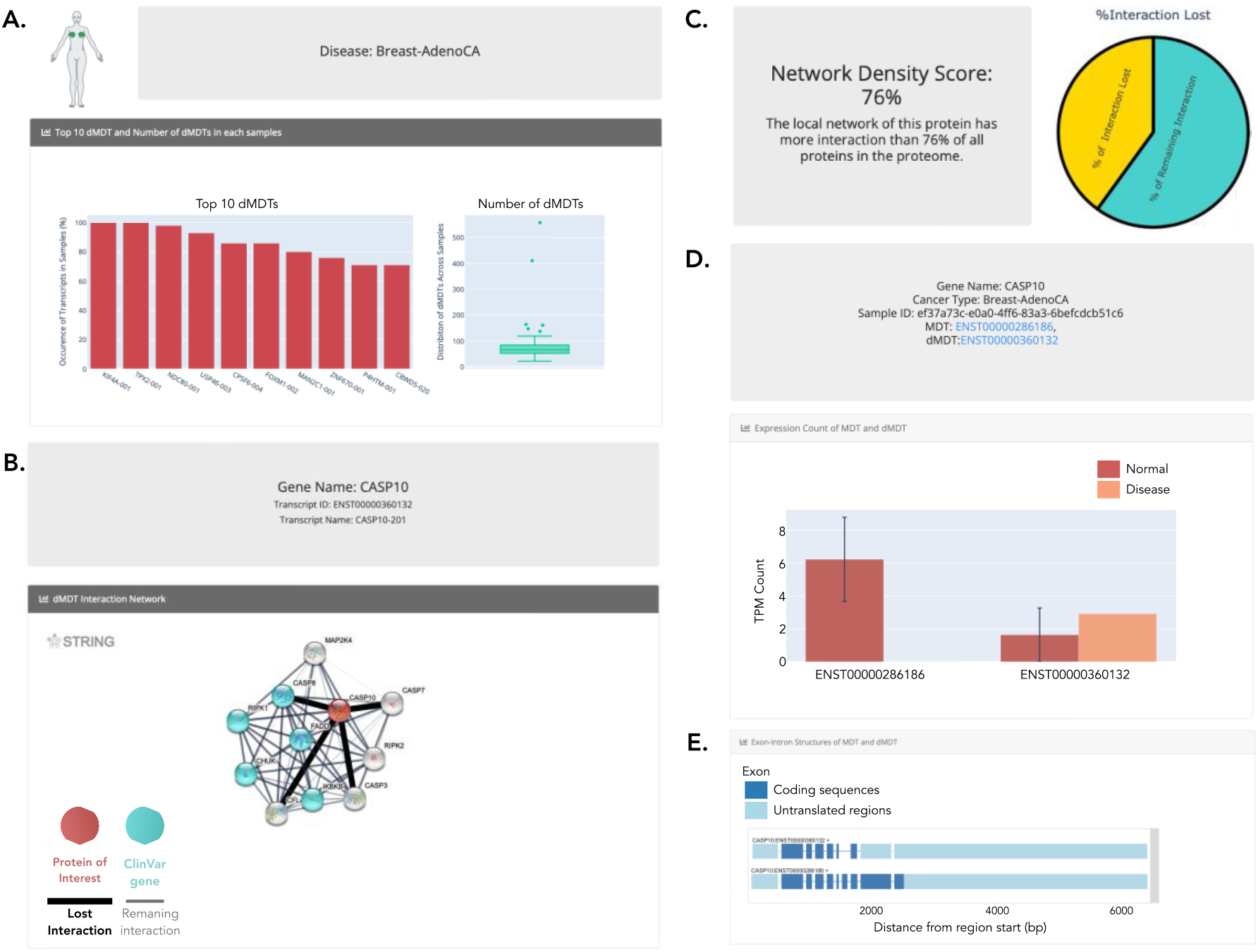
Exemplary case: **A.** Disease page; Top 10 dMDTs, which are most frequently observed in Breast-Adenocarcinoma samples. **B.** Isoform page; Interaction network of the isoform CASP10-201. Red sphere shows the protein of interest, spheres in turquoise highlight ClinVar genes. Black line represents the lost interaction from a dMDT, while gray lines represent the remaining interactions. **C.** Network density score and the percentage of interaction loss of the CASP10-201. **D.** Sample page; The expression level of MDT and dMDT in normal and disease sample. **E.** Exon-intron structure of MDT and dMDT in a specific sample.

CanIsoNet also provides disease-specific MDTs (dsMDT). dsMDT represent isoforms that are found as dMDT only in a single disease making them potential diagnostic biomarker candidates. The datasets used in CanIsoNet can be browsed and downloaded via dedicated drop-down menus or the download page. CanIsoNet is the first isoform-specific interaction network webserver for disease-specific isoforms. With its user-friendly interface as well as the rich annotations, CanIsoNet supports the discovery of functional and pathogenic isoform switching events. Upon request, new datasets can be uploaded to the CanIsoNet database.

## 4 Use Case

Opioid-induced hyperalgesia (OIH) is caused by enhanced pain response triggered by periodic administration of opioids. Understanding the molecular mechanism behind OIH would help to develop new treatments and relieve patients with chronic pain from permanent opioid usage (Ingram, 2022). Using the study GSE126662 in the NCBI GEO database, we have computed dMDT in OIH and uploaded the results to CanIsoNet for further investigation (see Supplementary File 1 for Method, Supplementary Table 1 for all detected dMDT). To access OIH specific dMDT data, select “Opioid-induced hyperalgesia” from the disease type table on the main page. This will direct load the OIH disease-specific page showing a barplot with the top 10 most frequent dMDT in the cohort, a boxplot highlighting the number of dMDT in each sample and two tables. The first table lists all switching events in OIH while the second table lists the total number of dMDT per sample that are visualized in the boxplot. Using the first table, the user can search for a specific isoform in the disease. For example, we observed a switching event in the gene *Irak3.* Irak3 is known as Interleukin-1 receptor-associated kinase 3, which is part of the Interleukin 1 signalling pathway (Jain *et al.*, 2014). Interestingly, disruption of the Interleukin-1 signalling pathway under opioid treatment are known to cause analgesic tolerance or OIH (Vanderwall and Milligan, 2019). Clicking on the dMDT transcript ID ENSMUST00000145665, opens a new website dedicated for visualizing information on the dMDT Irak3-002. The user can check the STRING interaction network and observe lost interactions due to domain-domain interaction losses. For Irak3, we detected an interaction loss to Myd88, which may trigger the hyperalgesia progression (Angst and Clark, 2006). Furthermore, a table is shown listing all lost interactions, the network density score at the location of the Irak3 gene and pie charts visualizing the percentage of lost interactions for Irak3 and the disease types. The latter information is also presented in a table, which holds links to samples in which the dMDT was observed. Clicking on the SRR8584386 sample ID, loads the sample specific page of CanIsoNet, which shows a barplot of the expression values of the dMDT in normal and in the disease sample. An exon-intron gene structure highlights differences in the exons and introns structures between the dMDT and MDT in the normal sample.

## Contribution

TK developed CanIsoNet webserver, TK, DS, CvM, HM, AK contributed to the webserver design, and CCS analysed OIH RNA-Seq samples. All authors contributed to writing the manuscript. AK had a role in every stage of the project.

## Funding

This project was funded by Krebsliga Zürich.

## Conflict of Interest

none declared.

## Notes

https://github.com/kahramanlab/CanIsoNet_Web

### Competing Interest Statement

The authors have declared no competing interest.

### Summary of Updates

- The functionality of web server has been extended to include other diseases. - The web server interface has been updated to make it more user-friendly. - An 'Use Case' has been added highlighting isoform switch events in Opioid-induced hyperalgesia. The following sections in the manuscript have been revised accordingly: - Abstract and Introduction part. - Result section has been updated with new disease results. - Figure 1 shows new web server interface. - A new author has been added. - Supplementary Files have been added.

## References

Angst, M.S. and Clark, J.D. (2006) Opioid-induced hyperalgesia: a qualitative systematic review. Anesthesiology, 104, 570–587.

Climente-González, H. et al. (2017) The Functional Impact of Alternative Splicing in Cancer. Cell Reports, 20, 2215–2226.

Ingram, S.L. (2022) Toward understanding the opioid paradox: cellular mechanisms of opioid-induced hyperalgesia. Neuropsychopharmacol., 47, 427–428.

Jain, A. et al. (2014) IL-1 Receptor-Associated Kinase Signaling and Its Role in Inflammation, Cancer Progression, and Therapy Resistance. Front. Immunol., 5.

Kahraman, A. et al. (2020) Pathogenic impact of transcript isoform switching in 1,209 cancer samples covering 27 cancer types using an isoform-specific interaction network. Sci Rep, 10, 14453.

Kaur Alasoo (2020) wiggleplotr: Make read coverage plots from BigWig files. R package version 1.14.0.

Landrum, M.J. et al. (2014) ClinVar: public archive of relationships among sequence variation and human phenotype. Nucl. Acids Res., 42, D980–D985.

Louadi, Z. et al. (2021) DIGGER: exploring the functional role of alternative splicing in protein interactions. Nucleic Acids Research, 49, D309–D318.

Maag, J.L.V. (2018) gganatogram: An R package for modular visualisation of anatograms and tissues based on ggplot2. F1000Res, 7, 1576.

Szklarczyk, D. et al. (2015) STRING v10: protein–protein interaction networks, integrated over the tree of life. Nucleic Acids Research, 43, D447–D452.

Vanderwall, A.G. and Milligan, E.D. (2019) Cytokines in Pain: Harnessing Endogenous Anti-Inflammatory Signaling for Improved Pain Management. Front. Immunol., 10, 3009.

Vitting-Seerup, K. and Sandelin, A. (2019) IsoformSwitchAnalyzeR: analysis of changes in genome-wide patterns of alternative splicing and its functional consequences. Bioinformatics, 35, 4469–4471.

Vitting-Seerup, K. and Sandelin, A. (2017) The Landscape of Isoform Switches in Human Cancers. Mol Cancer Res, 15, 1206–1220.

