## Supplementary File 1 for "CanIsoNet: A Database to Study the Functional Impact of Isoform Switching Events in Diseases"

[1] Institute of Molecular Life Sciences, University of Zurich, Zurich, Switzerland [2]Department of Pathology and Molecular Pathology, University Hospital Zurich, Zurich, Switzerland [3]Swiss Institute of Bioinformatics, Lausanne, Switzerland, [4]Computational Science and Engineering Department, Informatics Institute, Istanbul Technical University, Sariyer, Istanbul, 34467, Turkey, [5]Faculty of Medicine, University of Zurich, Zurich, Switzerland, [6]School for Life Sciences, Institute for Chemistry and Bioanalytics, University of Applied Sciences Northwestern Switzerland, Muttenz, Switzerland.

*corresponding author

### Supplementary Method

Here, most dominant transcript switching events for OIH were computed with the CanIsoNet software pipeline (Kahraman *et al.*, 2020) on mouse RNA-seq data obtained from the trigeminal ganglia (TG) which is a gathering location for sensory stimuli such as pain coming from face and the head regions (Gambeta *et al.*, 2020). (NCBI GEO: GSE126662). The RNA-seq data (100nt paired-end, RIN $\geq$ 7.5) was originally generated to evaluate the impact of morphine intake (0.9% in saline solution vs only saline solution) on the trigeminal ganglia and the nucleus accumbens in mice. The quality of the RNAseq data was checked using FastQC (Andrews, 2010) requiring a mean PHRED score >30 for each read. Reads were pseudo-aligned to the Ensembl Mouse database (GRCm38.p6) using Kallisto (Bray *et al.*, 2016). Transcripts Per Million (TPM) counts of identical cDNA sequences were merged and only the one with the longest protein isoform sequence were kept. TPM values < 2 were set to 0 given that 99% of olfactory receptor transcripts had a TPM value of < 2. A protein-protein interaction network for mus musculus was downloaded from the STRING database (v.11.0) (Szklarczyk *et al.*, 2015). An isoform-specific protein-protein interaction network was created as described in (Kahraman *et al.*, 2020). PFAM domain interactions from 3did (v.2020_01) (Mosca *et al.*, 2014) were integrated with STRING interactions and alternatively spliced protein isoform sequences from the Ensembl database (GRCm38.p6). To identify Most Dominant Transcript (MDT) switches between morphine treated and untreated mice, for each gene a MDT was identified if its expression was 1.5x higher than the 2nd most expressed transcript. Different from the original algorithm of CanIsoNet, we allowed 20% of untreated samples to share the same MDT with treated samples and relaxed the multiple testing q-value to < 0.1. In total we could identify 28 switch events (please see Supplementary Table S1 or https://www.caniso.net/Disease?disease=OIH).

### REFERENCES


Andrews,S. (2010) FastQC: A Quality Control Tool for High Throughput Sequence Data [Online]. Available online at: http://www.bioinformatics.babraham.ac.uk/projects/fastqc/.

Bray,N.L. *et al.* (2016) Near-optimal probabilistic RNA-seq quantification. *Nat Biotechnol*, **34**, 525–527.

Gambeta,E. *et al.* (2020) Trigeminal neuralgia: An overview from pathophysiology to pharmacological treatments. *Mol Pain*, **16**, 174480692090189.

Kahraman,A. *et al.* (2020) Pathogenic impact of transcript isoform switching in 1,209 cancer samples covering 27 cancer types using an isoform-specific interaction network. *Sci Rep*, **10**, 14453.

Mosca,R. *et al.* (2014) 3did: a catalog of domain-based interactions of known three-dimensional structure. *Nucl. Acids Res.*, **42**, D374–D379.

Szklarczyk,D. *et al.* (2015) STRING v10: protein–protein interaction networks, integrated over the tree of life. *Nucleic Acids Research*, **43**, D447–D452.
